# Genetic silencing of K_Ca_3.1 inhibits atherosclerosis in ApoE null mice

**DOI:** 10.1101/2024.12.11.628017

**Authors:** P. Alam, D.L. Tharp, H.J. Bowles, L. Grisanti, H. Bui, S.B. Bender, D.K. Bowles

## Abstract

Increased expression of K_Ca_3.1 has been found in vascular smooth muscle (SMC), macrophages, and T cells in atherosclerotic lesions from humans and mice. Proliferating SMC cells increase the expression of K_Ca_3.1, such that it becomes a dominant K^+^ channel and contributes to SMC cell migration. The efficacy of pharmacological inhibition of K_Ca_3.1 in limiting atherosclerosis progression has been demonstrated in mice and pigs, however direct, loss-of-function, i.e. gene silencing, studies are absent. To investigate the role of K_Ca_3.1, we used CRISPR/Cas9 to generate K_Ca_3.1^-/-^Apoe^-/-^ (DKO) mice and assessed lesion development in the brachiocephalic artery (BCA) of DKO versus Apoe^-/-^ mice on a Western diet for 3 months. Notably, the loss of K_Ca_3.1 did not affect serum total cholesterol or body weight. In BCAs of DKO mice, lesion size (0.036 mm² vs. 0.118 mm², p<0.05) and relative stenosis (13.9% vs. 43.0%, p<0.05) were reduced by 70% compared to Apoe^-/-^ mice, with no effect on medial or lumen area. Additionally, DKO mice exhibited a significant reduction in macrophage content within atherosclerotic plaques compared to Apoe^-/-^ mice, independent of sex. *In vitro* migration assays further showed a significant reduction in migration of bone marrow-derived macrophages (BMDMs) from DKO mice compared to those from Apoe^-/-^ mice. Furthermore, *in vitro* experiments using rat aortic smooth muscle cells (RAOSMCs) revealed significant inhibition of PDGF-BB-induced MCP1/Ccl21 expression upon K_Ca_3.1 inhibition, while activation of K_Ca_3.1 further enhanced MCP1/Ccl21 expression. Both *in vivo* and *in vitro* analyses showed that silencing K_Ca_3.1 and sex had no significant effect on the collagen content of plaque. RNAseq analysis of BCA samples from DKO and Apoe^-/-^ mice revealed PPAR-dependent signaling as a potential key mediator of the reduction in atherosclerosis due to K_Ca_3.1 silencing. Overall, this study provides the first genetic evidence that K_Ca_3.1 is a critical regulator of atherosclerotic lesion development and composition and provides novel mechanistic insight into the link between K_Ca_3.1 and atherosclerosis.

## Introduction

Despite lipid-lowering and emerging anti-inflammatory agents, atherosclerosis remains the leading cause of death in both men and women in the United States ^1,2^. Over 20 million Americans > 20 years of age have coronary heart disease (CHD), and each year ∼635,000 Americans have a new coronary attack, and ∼300,000 have a recurrent attack^3^. Atherosclerosis is a chronic, inflammatory, and proliferative disease that develops over decades, involving multiple cell types, including endothelial cells, smooth muscle cells (SMCs), fibroblasts, macrophages, T-cells, B-cells, and platelets. The intermediate-conductance Ca^2+^ -activated K^+^ channel (K_Ca_3.1) is expressed in all of these cell types and plays a crucial role in T-cell, B-cell, fibroblast, and SMC proliferation, as well as the migration of SMCs, macrophages, and platelet coagulation^4–8^, leading to the consideration of K_Ca_3.1 modulators as potential therapies for vascular disease^9,10^.

Increased expression of K_Ca_3.1 has been found in atherosclerotic lesions from humans and mice^7,11^ and several studies have shown systemic delivery of K_Ca_3.1 inhibitors can attenuate atherosclerosis lesion development in mice^7,11–13^. Synthetic, proliferating SMC cells increase expression of K_Ca_3.1, such that it becomes the dominant K^+^ channel ^8,14^. Although the cell type-specific relative contribution of K_Ca_3.1 activation during atherosclerosis has not been determined, its upregulation has been observed in neointimal SMCs in balloon-injured rat carotid arteries^7,15^ as well as atherosclerotic lesions in mice and humans^7^. In addition, we have shown that acute administration of the K_Ca_3.1 inhibitor, TRAM-34, during coronary angioplasty in a swine model of coronary restenosis can inhibit lesion development, predominantly by targeting SMC proliferation ^5^.

Migration of SMC from the media to the intima is a major contributor to both restenosis and atherosclerosis and we^5,8^ and others ^6,7^ have shown that K_Ca_3.1 is critical for SMC migration. Together, these studies support the hypothesis that the upregulation of K_Ca_3.1 is a major contributor to SMC migration and proliferation during atherosclerosis and restenosis. In addition to potential effects on SMC, K_Ca_3.1 inhibitors^7,11^ reduce lesion macrophage content in Apoe^-/-^ mice. In macrophages, K_Ca_3.1 regulates migration, M1/M2 polarization, respiratory burst, and pathogen killing^16^. Pharmacological blockade of K_Ca_3.1 inhibits M1 polarization and increases M2/M1 ratio in advanced plaques^11^. In addition to the direct role of macrophage K_Ca_3.1 in activation, K_Ca_3.1 may also regulate interactions between SMC and macrophages by modulating the inflammatory phenotype of SMCs^10^. The “inflammatory state” of SMCs in the lesion cap is crucial, as these cells secrete chemokines such as MCP1/Ccl21^17^, which recruit monocytes/macrophages that degrade the fibrous cap and contribute to the plaque instability^18,19^.

In summary, while there is substantial evidence supporting the role of K_Ca_3.1 in SMC and macrophage function in atherosclerosis, the evidence to date is derived from *in vitro* studies on selected cell types or *in vivo* pharmacological investigations. The purpose of the current study was to provide the first genetic silencing of K_Ca_3.1 in the context of atherosclerosis development *in vivo*.

## Material and Methods

### Ethics Statement

Experimental protocols were approved by the University of Missouri Animal Care and Use Committee and in accordance with the “Principles for the Utilization and Care of Vertebrate Animals used in testing, Research and Training.”

### Generation of Apoe^-/-^Kcnn4^-/-^ (DKO) mice

To firmly establish a genetic causal link between K_Ca_3.1 and atherosclerosis, we developed a constitutive, double knockout mouse silencing both K_Ca_3.1 and ApoE (DKO). All mouse models were generated on the C57BL/6 background. We used a CRISPR/Cas9 approach to create a DKO in conjunction with the MU Animal Modeling Core. sgRNA sequences were designed and produced (Sage Research Labs, St. Louis, MO). gRNA targeting exon 4 of the Kcnn4 gene (gRNA/*PAM site*; tcagcgccacgcgccagtcc*agg*) resulted in an 11 bp deletion (Δ887-907) and frameshift mutation, while a dual gRNA targeting exon 3 of Apoe (gRNA/*PAM site*: upstream, gtaactcaagctggcttcga*agg*; downstream, ggtaatcccagaagcggttc*agg*) resulted in deletion of ApoE (Figure 1a). Because Apoe and Kcnn4 both reside on chromosome 7, it was not possible to generate wild-type littermates from Apoe^-/-^Kcnn4^+/+^ crosses for controls, therefore Apoe^-/-^ mice (B6.129P2-Apoe^tm1Unc^/J; Jackson Laboratory, Bar Harbor, MN) were bred for comparison. Male and female mice were fed ad libitum with standard chow diet until 8-10 weeks of age then switched to a Western diet (0.2% cholesterol, 42% kcal from fat; TD.88137, Teklad Diets) for 16 weeks. Mice were anesthetized, and either perfusion-fixed via the left ventricle at 80 mmHg with 10% neutral-buffered formalin for histological analysis, or the brachiocephalic artery (BCA) was carefully dissected and rapidly frozen in liquid nitrogen for molecular analysis. The BCA from perfusion-fixed animals was then dissected and stored in formalin until it was embedded in paraffin for subsequent histology.

**Figure 1.**
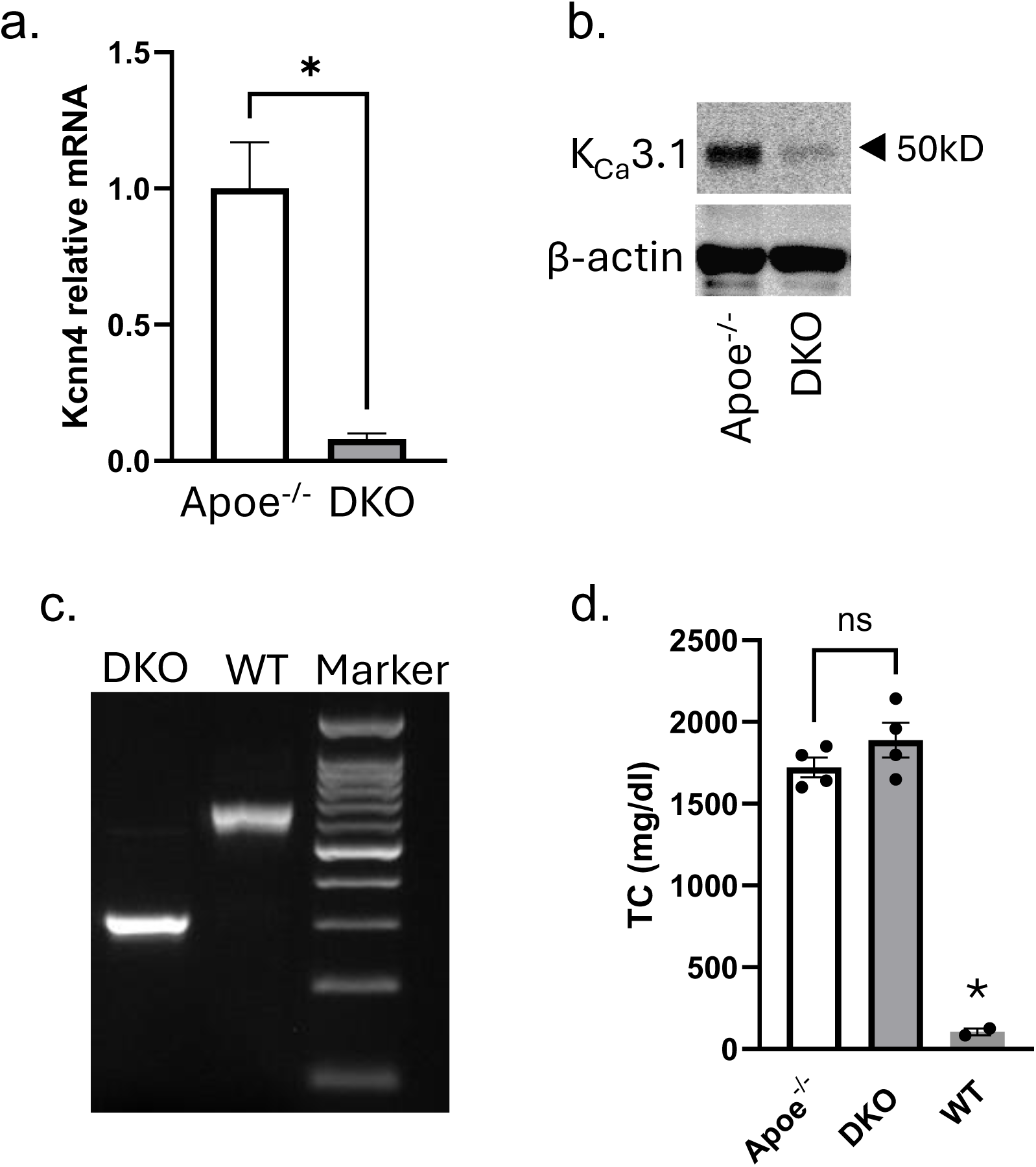
K_Ca_3.1 silencing in DKO mouse. Loss of K_Ca_3.1 mRNA (a) and protein (b) in DKO compared to Apoe^-/-^, c) PCR gel of Apoe showing 500 bp deletion in DKO compared to WT d) producing similar elevated levels of total plasma cholesterol in DKO and Apoe^-/-^ vs. WT. *p < 0.05.

### Immunohistochemistry

Immunohistochemistry was performed as previously described^5^. Sections were incubated with avidin–biotin two-step blocking solution (Vector SP-2001) to inhibit background staining and 3% hydrogen peroxide to inhibit endogenous peroxidase. Non-serum protein block (Dako X909) was then applied to inhibit non-specific protein binding. Sections were incubated at 4°C overnight with primary antibodies SMαA (Dako, #M0851), or CD68 (Invitrogen, #MA513324). After washing, sections were incubated with biotinylated secondary antibody in phosphate-buffered saline containing 15 mM sodium azide and peroxidase-labeled streptavidin (Dako LSAB+ kit, peroxidase, K0690). Diaminobenzidine (DAB, Dako) was applied 5 minutes for visualization of the reaction product, sections were then counterstained with hematoxylin, dehydrated, and coverslipped. Images of the sections were obtained using an Olympus BX61 photomicroscope and Spot Insight Color camera (Diagnostic Instruments). The relative area and mean density of positive staining were determined for each section of interest utilizing ImagePro Plus (Media Cybernetics).

### Morphology

Serial sections were obtained from the BCA and stained using Verhoeff-Van Gieson (VVG) and picrosirius red. Morphometric measurements were obtained on spatially calibrated images of VVG stained sections using standard planimetry and NIH Image J software (Bethesda, MD). Intimal area was defined as the area between the internal elastic lamina (IEL) and the luminal border of the artery while the area between the IEL and external-elastic lamina (EEL) was referred to as the medial area. Intima-media ratio (I/M) was calculated as the ratio of intimal-over medial-areas. Percent stenosis was defined as intimal area/IEL area. Relative intimal collagen content was determined from Picrosirius red images captured under polarized light. Necrotic core (NC) was defined as acellular areas within a lesion that were negative for VVG staining (i.e., extracellular matrix was lacking with total or almost complete loss of collagen). Boundary lines were delineated around those regions and the area was measured by image analysis software as described above. Based on the criterion by Seimon et. al. ^20^, areas <3,000 μm^2^ were not counted as they likely do not constitute substantial areas of necrosis. Relative NC area was calculated by dividing total NC area by total lesion area.

### Blood analysis

Total cholesterol content was determined by Comparative Clinical Pathology Services, LLC (Columbia, MO) from serum collected the day of euthanasia.

### Chemotaxis

Bone marrow-derived macrophages (BMDM) from Apoe^-/-^ and DKO mice were obtained as described previously^21^. BMDM were plated at 30,000-40,000 cells per well in the upper chamber of an 8µm pore 24-well chemotaxis chamber (Millipore). The following solutions were placed in the lower chamber (diluted in serum-free media): vehicle or MCP1/Ccl2 (10 ng/mL). The chamber was placed at 37°C for 4 hours. Cells from the upper chamber were removed, and the filters were stained using the Diff-Quik Staining Kit (Fisher scientific). The migrated cells in a 40X field were manually counted.

### Rat aortic smooth muscle cell (RASMC) culture

RASMC were obtained from Cell Application (catalog # R354-05a). RASMCs were grown to post-confluence and serum-starved for six days. Following starvation, cells were treated with TRAM-34 or SKA-31, either alone or in combination with PDGF-BB. Twenty-four hours post-treatment, cells were harvested for qRT-PCR analysis of K_Ca_3.1, Ccl2, and Col1a1. For siRNA-mediated inhibition of K_Ca_3.1, RASMCs were transfected with 150 nM of siK_Ca_3.1, and a scrambled siRNA (siCnt) was used as a control. Twenty-four hours post-transfection, cells were serum-starved for 24 hours to maximize the expression of smooth muscle differentiation markers^8^. Cells were then treated with either vehicle or Platelet-Derived Growth Factor (PDGF)-BB (20 ng/mL) for an additional 24 hours, after which RNA was isolated for qRT-PCR-based gene expression analysis.

### RNA isolation

RNA isolation from BCA and aorta was performed by using the RNeasy Plus Mini Kit (Quagen Catalog # 74136) following the standard protocol. Briefly, sample lysis buffer was added to the BCA and aorta tissue samples. Then, BigPrep Lysing Matrix D was added to each sample, and the tubes were placed in a FastPrep homogenizer. The samples were homogenized for 20-30 seconds at maximum speed (6.0 m/s) to ensure complete tissue disruption. The process was repeated as needed to achieve a homogeneous sample for further analysis. After homogenization, the samples were passed through a gDNA column to remove genomic DNA. The flow-through was then mixed with an equal volume of 70% ethanol. The mixture was subsequently applied to an RNA elution column, followed by washing steps according to the kit’s instructions. RNA was then eluted, and the concentration and quality of the RNA were assessed using a Nanodrop spectrophotometer. The same RNA isolation protocol was followed for the *in vitro* cell culture experiments, with the exception of the FastPrep homogenization step.

### Quantitative reverse-transcriptase PCR (qRT-PCR)

Quantitative RT-PCR was performed as previously described^8,22,23^. Samples were quick frozen in liquid nitrogen and stored at −80°C until processed. Cultured cells were frozen in TRIzol solution. Total RNA was isolated according to the TRIzol published protocol. cDNA was transcribed from total RNA using High-Capacity cDNA reverse transcriptase kit (Applied Biosystems, Catalog# 4368814). A minus reverse transcriptase reaction was performed to ensure no genomic DNA contamination. Quantitative RT-PCR was performed on a MyiQ iCycler (Bio-Rad, model 170-9770). Each 20uL reaction contained 1X Syber Green Master Mix (Bio-Rad), 0.8µM forward and reverse primers, and 1µg of cDNA. Each reaction was initiated by a 95°C hold for 3 minutes to activate heat stable Taq polymerase, and reaction conditions were optimized for each set of primers. Target gene expression was normalized to 18S ribosomal RNA or Ubiquitin C (UBC) using the 2^-ΔΔCT^ method ^24^.

### Transcriptomic analysis

For transcriptomic analysis, 1 or 2 pooled BCAs were snap-frozen to create samples from both groups (n=10-11 per group). Sequencing was conducted by the MU DNA Core. To generate Illumina Stranded mRNA libraries, the poly-A containing mRNA was purified from total RNA, RNA was fragmented, double-stranded cDNA was generated from fragmented RNA, and the unique dual index containing adapters were ligated to the fragment ends. All 40 libraries generated were sequenced together on an Illumina NovaSeq flow cell using paired-end read, 100 base read length sequencing. Total yield was ∼2 billion reads providing ∼50 million read pairs per sample. Analysis and visualization of the NGS sequence data was performed by the MU Informatics Research Core Facility (IRCF).

RNA-Seq data were processed and analyzed as previously described^25^. Briefly, latent Illumina adapter sequence was identified and removed from input 100-mer RNA-Seq data using Cutadapt. Subsequently, input RNA-Seq reads were trimmed and filtered to remove low quality nucleotide calls and whole reads, respectively, using the Fastx-Toolkit. To generate the final set of quality-controlled RNA-Seq reads, foreign or undesirable sequences were removed by similarity matching to the Phi-X genome (NC_001422.1), the relevant ribosomal RNA genes as downloaded from the National Center for Biotechnology Information, or repeat elements in RepBase, using Bowtie. The remaining RNA-seq reads were aligned to the Mus musculus genome assembly GRCm39 using the Bioconductor package Rsubread with default settings for paired reads. Rsubread uses a “seed-and-vote” mapping paradigm which allows for quick mapping of subreads for gene expression analysis. This provided the initial gene expression estimates which were annotated according to their Entrez ID number using the Bioconductor database org.Mm.eg.db. These initial expression estimates were then normalized and sorted into a comparison matrix based on genotype using the Bioconductor package DESeq2. The two comparison groups, Apoe-/- and DKO, were then analyzed according to their differential gene expressions with DKO used as the treatment group to be compared against the Apoe-/- expressions. Outlier samples were identified through the use of Principal Component Analysis (PCA) and Counts Per Million plots. Normalization and differential gene expression (DGE) analysis was then rerun after the exclusion of aforementioned outliers. The final results were written into an excel file and visualized using PCA and volcano plots, highlighting genes with a log2(fold change) >2 and adjusted p-values <0.05. Genes with the lowest adjusted p-values were analyzed using DAVID (david.abcc.ncifcrf.gov)^26^ and IPA (Ingenuity Systems) as previous^27–32^.

### Statistical analysis

All data are presented as mean ± SE. One-way or two-way ANOVA was used for all group comparisons as appropriate, and significance was defined as p ≤ 0.05. The Student’s *t*-test was applied for comparisons between two groups. Statistical cutoffs for differentially expressed genes were −1.0 ≤ log^2^FC ≥ 1.0 change in expression with an adjusted p < 0.05.

## Results

### Apoe^-/-^Kcnn4^-/-^ (DKO) mice

In DKO mice, we confirmed a significant reduction in K_Ca_3.1 mRNA (Fig. 1a) and protein (Fig. 1b) expression compared to Apoe^-/-^ mice. Genotyping of the Apoe allele showed a 500 bp deletion in the DKO mice, when compared to WT mice (Fig. 1c). Founder DKO mice also demonstrated a similar increase in plasma cholesterol levels as Apoe^-/-^ mice compared to wild-type controls. Importantly, no significant difference was observed in cholesterol levels between DKO and Apoe^-/-^ mice (Fig. 1d).

### Group characteristics

Group comparisons for body weight (BW) and total cholesterol are shown in Table 1. Overall, male mice were heavier than females across genotype. Male Apoe^-/-^ and DKO body weights were similar, while DKO female mice were heavier than Apoe^-/-^ females. While total cholesterol was lower in Apoe^-/-^ females compared to Apoe^-/-^ males, K_Ca_3.1 silencing had no effect within either sex.

**Table 1.**

|  | Male |  | Female |  |
| --- | --- | --- | --- | --- |
|  | Apoe<sup>-/-</sup> | DKO | Apoe<sup>-/-</sup> | DKO |
| BW (g) | 35.4 | 37.2 | 25.8<sup>#</sup> | 29.8<sup>*#</sup> |
| SE | ±1.3 | ±1.4 | ±0.7 | ±0.9 |
| N | 12 | 12 | 12 | 9 |
| TC (mg/dl) | 1670 | 1706 | 1303<sup>#</sup> | 1521 |
| SE | ±76 | ±124 | ±64 | ±75 |
| N | 12 | 12 | 12 | 9 |

\*p<0.05 vs. Apoe<sup>-/-</sup>; # vs. male

### Brachiocephalic artery morphometry

We examined atherosclerosis development in the brachiocephalic artery (BCA). As reported previously^33^, Apoe^-/-^ mice fed a Western diet developed complex atherosclerotic lesions consisting of a fibrous, cellular matrix overlying lipid cores (Fig. 2a). Although both neointimal area and relative stenosis were lower in female compared to male mice in both Apo^-/-^ and DKO mice, DKO mice demonstrated significantly reduced neointimal size (NI; Fig. 2a, c) and relative stenosis (Fig. 2b, d) in both sexes.

**Figure 2.**
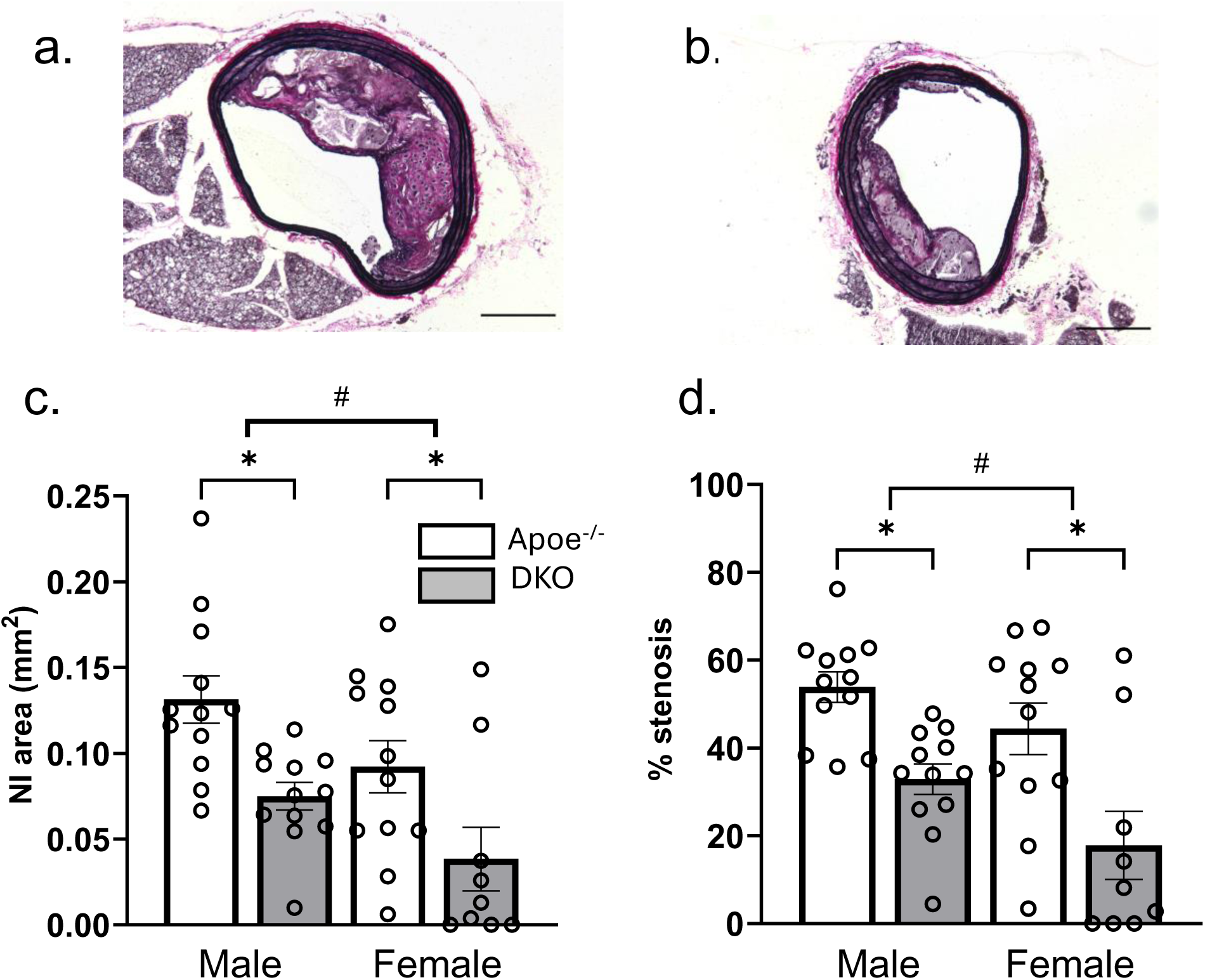
K_Ca_3.1 silencing reduces atherosclerotic plaque size. Representative VVG stained BCA sections Apoe^-/-^ (a) and DKO (b) mice. Scale bar = 200 µm. Morphometric data showing (c) neointimal (NI) areas and (d) % stenosis in Apoe^-/-^ and DKO males and females. *p<0.05, #p<0.05 main effect of male vs. female. While females had overall smaller NI and % stenosis vs. males in both groups, silencing K_Ca_3.1 significantly reduced both NI area and % stenosis in both sexes. n=12 and 12 for Apoe^-/-^ male and female, resp., and n=12 and 9 DKO male and female, resp.

### Silencing K_Ca_3.1 alters atherosclerotic plaque composition

We and others have previously shown that inhibiting K_Ca_3.1 reduces smooth muscle cell proliferation and migration and macrophage activation and migration^7,8,11^. Consistent with these findings, both smooth muscle cell (Fig. 3a-c) and macrophage (Fig. 3d-f) content were significantly reduced in lesions from DKO mice when compared to those from Apoe^-/-^ mice. It has been shown the majority of SMC within atherosclerotic lesions are of medial origin and migrate into the intima during lesion development^34–36^. Previous studies by us and others ^7,8,22^ have shown the K_Ca_3.1 inhibitor, TRAM-34, inhibits SMC migration *in vitro* and reduces SMC content in atherosclerotic plaques. The effect of K_Ca_3.1 silencing in reducing SMC content within the lesion suggests that K_Ca_3.1 activation is involved in medial-to-intimal SMC migration in atherosclerosis development. Additionally, our results show a significant reduction in the necrotic core size in DKO mice compared to Apoe^-/-^ mice (Fig. 4a-c). In contrast, K_Ca_3.1 silencing, and sex had no significant effect on the relative collagen content in the lesions (Fig. 4d-f). Overall, lesions in DKO animals were smaller, with reduced smooth muscle and macrophage content, and a dramatically diminished necrotic core. Together, these findings suggest that K_Ca_3.1 silencing leads to smaller lesions with features indicative of a more stable plaque.

**Figure 3.**
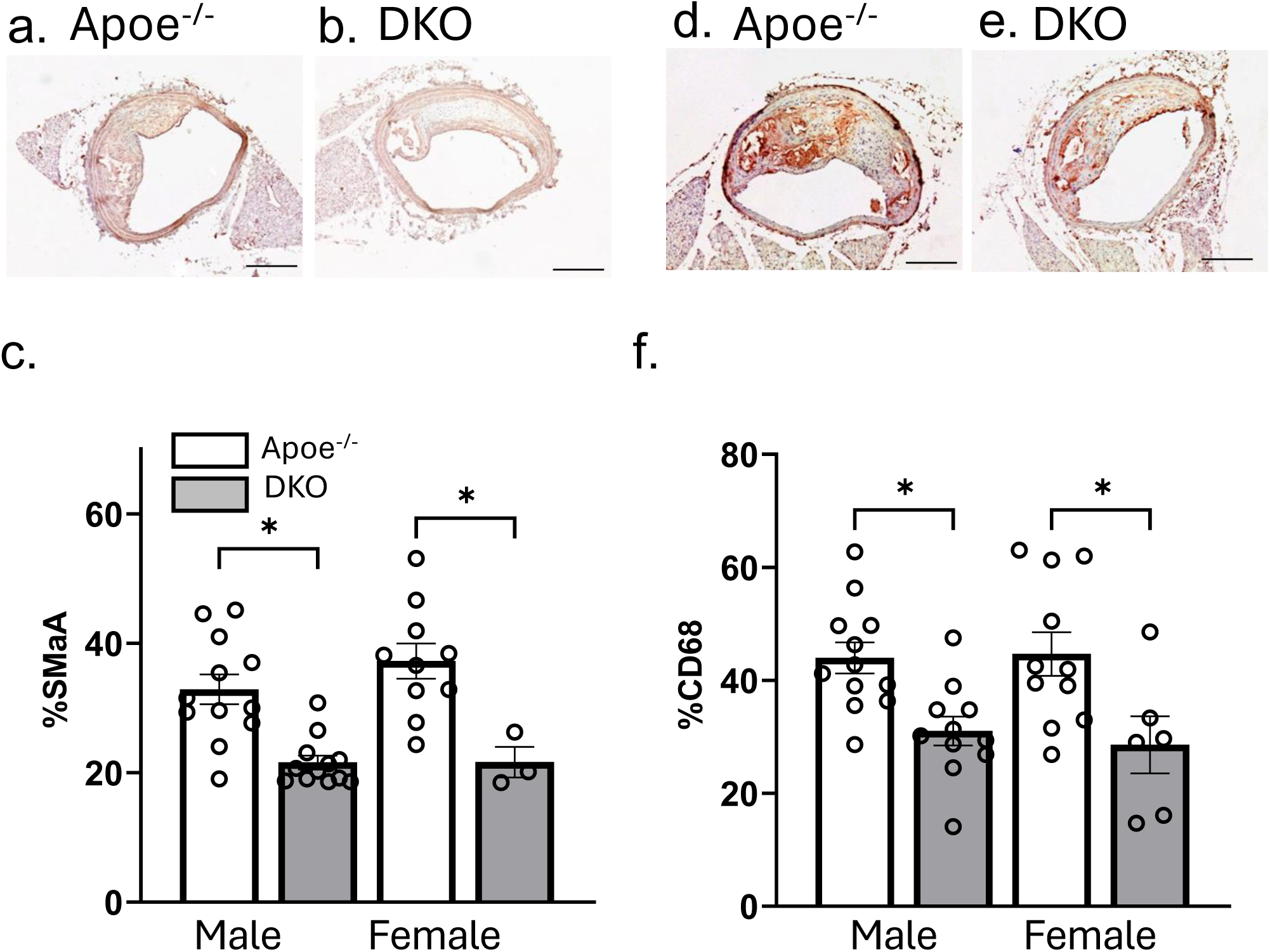
K_Ca_3.1 silencing reduces intimal smooth muscle (a-c) and macrophage (d-f) content. Representative BCA sections from Apoe^-/-^ (a) and DKO (b) mice probed with anti-smooth muscle alpha actin (SMαA) and (c) group data for relative SMαA positive intimal area presented as mean ± S.E, n=12 and 10 for Apoe^-/-^ male and female, resp., and n=12 and 3 DKO male and female, resp. Representative BCA sections from Apoe^-/-^ (d) and DKO (e) mice probed with CD68 and (f) group data for relative CD68 positive intimal area presented as mean ± S.E, n=12 and 11 for Apoe^-/-^ male and female, resp., and n=11 and 6 DKO male and female, resp. resp.*p<0.05.

**Figure 4.**
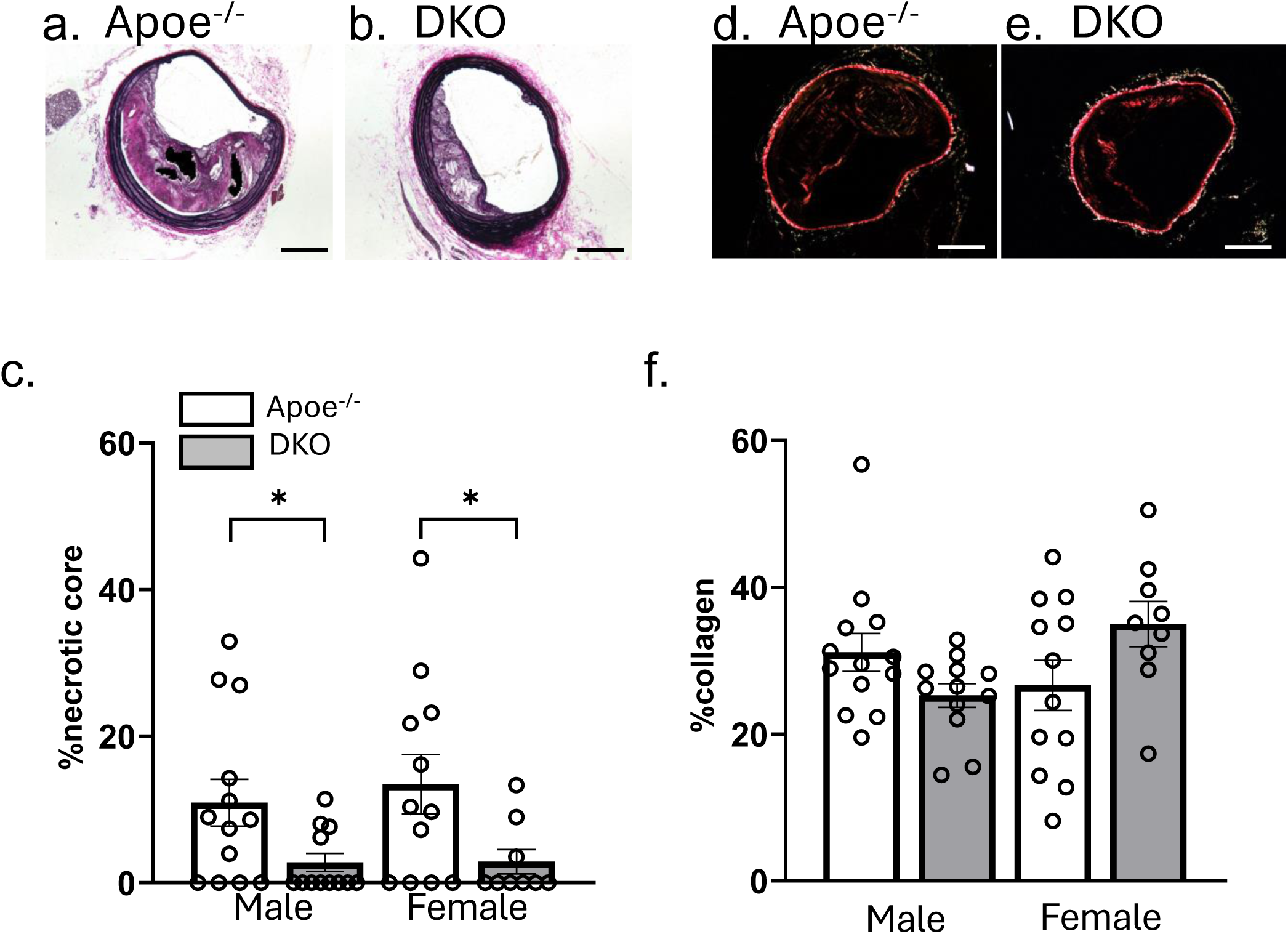
K_Ca_3.1 silencing reduces necrotic core size with no effect on relative intimal collagen content. Representative VVG stained BCA sections from Apoe^-/-^ (a) and DKO (b) with indicated necrotic core (black area) as defined in Methods and (c) group data for relative necrotic core area. Representative BCA sections from Apoe^-/-^ (d) and DKO (e) mice stained with picrosirius red under polarized light and (f) group data for relative positive intimal area. Data are presented as mean ± S.E, n=13 and 12 for Apoe^-/-^ male and female, resp., and n=12 and 9 DKO male and female, resp.

We performed *in vitro* studies to investigate potential mechanisms of reduced macrophage content in atherosclerotic lesions after K_Ca_3.1 silencing. Migration of BMDM from DKO mice was significantly reduced compared to Apoe^-/-^ in both male and female mice (Fig. 5), providing direct genetic evidence supporting previous findings demonstrating a role for K_Ca_3.1 in macrophage activation and infiltration^7,11,37^. In addition, given the potential for intimal SMCs to contribute to macrophage infiltration via expression of chemotactic factors, such as MCP1/Ccl2, we examined the role of K_Ca_3.1 in MCP1/Ccl2 expression in SMC (Figure 6). In rat aortic smooth muscle cells (RASMCs) K_Ca_3.1 activity was modulated using specific siRNA (siK_Ca_3.1), along with its pharmacological inhibition by TRAM-34 and activation by SKA-31. As previously, we observed a significant increase in K_Ca_3.1 expression after PDGF-BB treatment^8^, which was significantly inhibited following K_Ca_3.1 inhibition by either siK_Ca_3.1 or TRAM-34, and augmented by SKA-31 (Fig. 6a-c) confirming a positive feedback role of K_Ca_3.1 on self-expression^5,8^. Similarly, PDGF-BB-induced MCP1/Ccl2 expression was inhibited following K_Ca_3.1 inhibition by either siK_Ca_3.1 or TRAM-34 and augmented by SKA-31 (Fig. 6d-f). Furthermore, using rat aortic endothelial cells (RAOECs), we observed no effect of K_Ca_3.1 inhibition, either by siRNA (Fig. 6g) or the inhibitor TRAM34 (Fig. 6h), nor its activation by SKA31 (Fig. 6i), on the expression of Ccl2. This suggests that the observed changes in Ccl2 expression are likely derived from SMCs. In contrast, no significant effect was observed on Col1a1 expression following either PDGF-BB treatment or inhibition of K_Ca_3.1 (Supplementary Fig. 1a,b), however we did observe a main effect of K_Ca_3.1 activation by SKA-31 (Supplementary Fig. 1c). Thus, these studies confirm a putative multicellular contribution of K_Ca_3.1 to macrophage content of atherosclerotic lesions.

**Figure 5.**
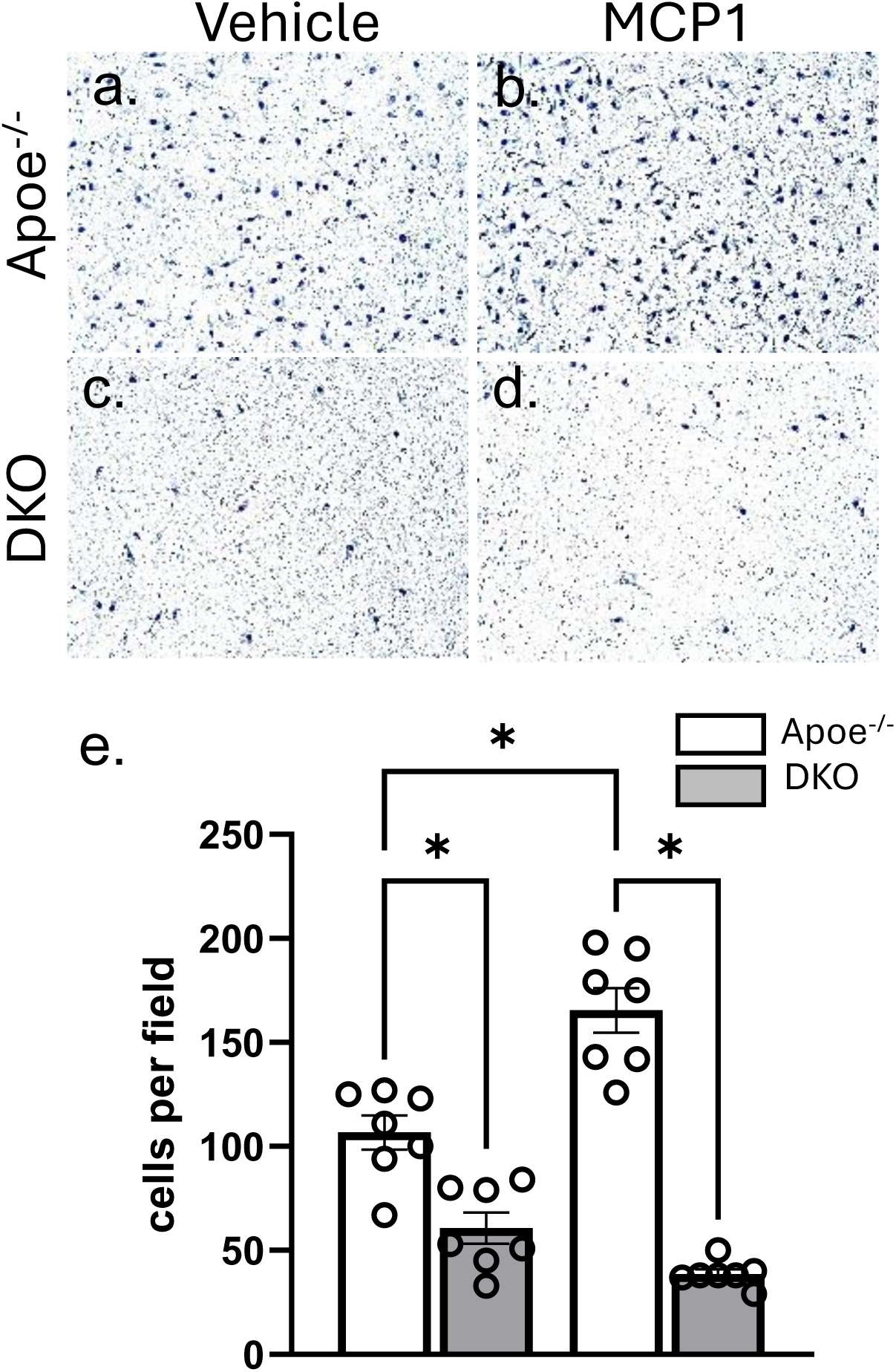
Genetic ablation of K_Ca_3.1 inhibits macrophage migration in vitro. Representative membranes from chemotaxis chambers showing migrated BMDM cells from Apoe^-/-^ (a,b) and DKO (c,d) in response to vehicle (a,c) or Ccl2/ MCP1 stimulation (b,d). Quantified group data (e) presented as mean ± S.E n=7 per group.

**Figure 6.**
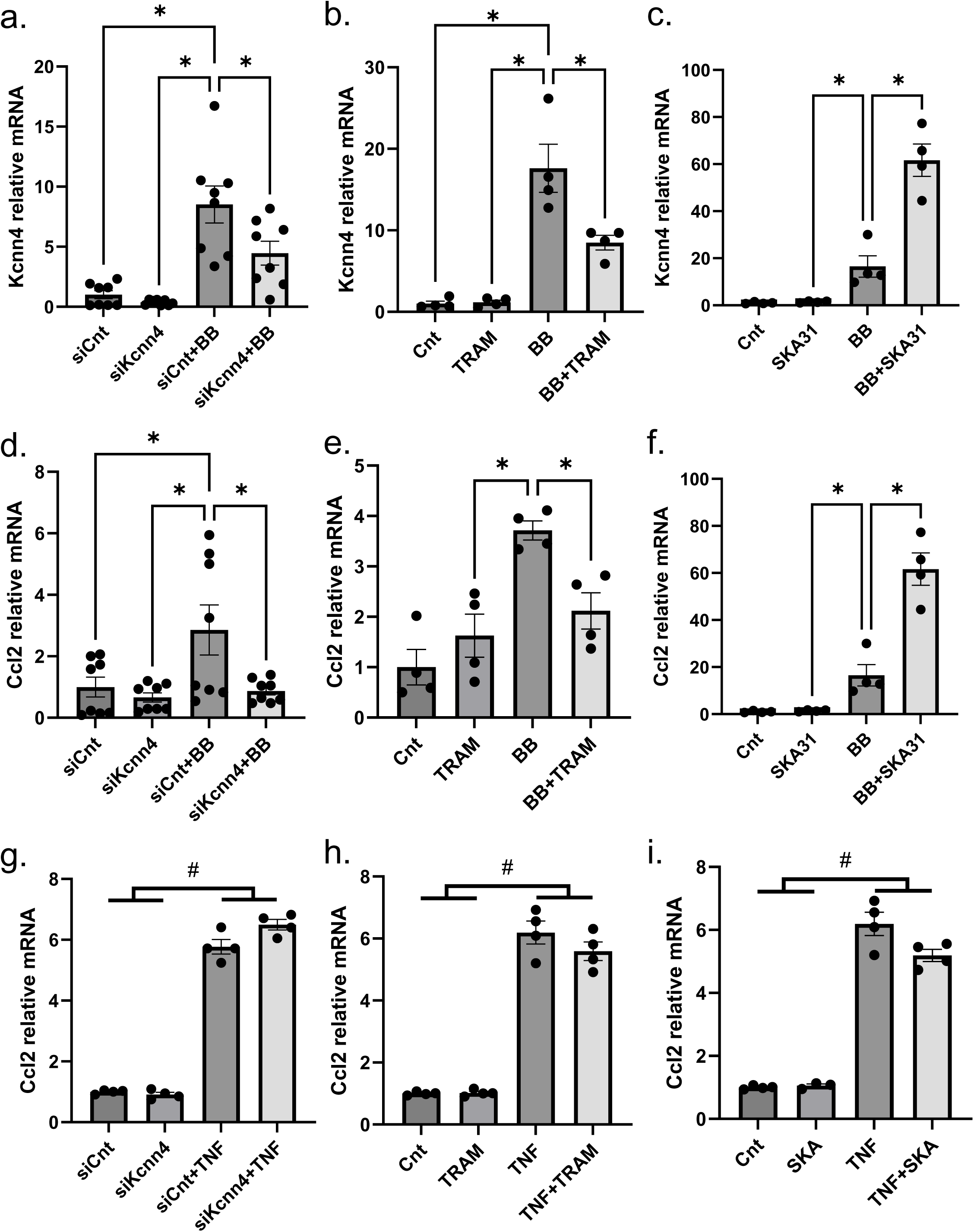
K_Ca_3.1 regulates MCP1/Ccl2 expression in smooth muscle, but not endothelial cells. RASMCs were stimulated with and without PDGF-BB (20 ng/ml) for 24 hours while K_Ca_3.1 was either inhibited by siRNA (a,d,g) or TRAM 34 (b,e,h) or, conversely, stimulated with the K_Ca_3.1 activator, SKA-31 (c,f,i). The PDGF-BB induced increase in both Kcnn4 and Ccl2 was inhibited by either genetic (a,d) or pharmacological inhibition (b,e) of K_Ca_3.1, while conversely, stimulation of K_Ca_3.1 greatly augmented the response (c,f). K_Ca_3.1 inhibition, or activation in RAOEC does not affect the Ccl2 expression (g-i). Data represented as mean±SE. the *p*-value ≤0.05 was considered statistically significant. * *p*-value ≤0.05. # *p*-value >0.05.

### Putative beneficial transcriptome changes in atherosclerotic lesions with K_Ca_3.1 silencing

In order to explore potential novel mechanisms underlying the beneficial impact of K_Ca_3.1 silencing, we performed bulk RNA sequencing of diseased BCA from both Apoe^-/-^ and DKO mice to identify unique transcriptomic signatures induced by K_Ca_3.1 silencing in atherosclerosis. Analysis identified 449 differentially expressed genes (DEG) significantly altered (152 downregulated and 297 upregulated, Figure 7a). DEGs were used to identify key pathways using gene ontology (GO) enrichment analysis and directional gene changes in IPA. The GO analysis using Metascape (https://metascape.org) revealed significant activation of the respiratory electron transport chain, PPAR signaling pathways, and metabolic pathways (Supplementary Fig. 2a). Moreover, regulatory factor analysis demonstrated a predominant role of PPARs (Supplementary Fig. 2b). Similarly, IPA identified the top four statistically enriched pathways as mitochondrial, i.e. decreased mitochondrial dysfunction, increase respiratory electron transport, increases oxidative phosphorylation and increases mitochondrial fatty acid oxidation (Fig. 7b). Comparison analysis within IPA aligned DKO with great similarity to the top enriched pathways identified from transcriptomic analysis of rosiglitazone treatment (Fig. 7b). DEGs within the PPAR signaling pathway, included significant upregulation of PPARα, PPARγ and PPARGC1A (Fig. 7c). Accordingly, IPA identified the PPAR agonist, rosiglitazone, as the top upstream regulator activated with 45 of 52 predicted downstream genes activated (Supplementary Table 1). Overall, transcriptomic analysis identified PPAR pathway activation as a top putative mechanism underlying the effects of K_Ca_3.1 silencing on atherosclerosis. Examination of top DEG pathways downstream of K_Ca_3.1 inhibition with cellular processes known to be involved in atherosclerosis reveals a novel model of the mechanisms underlying K_Ca_3.1 silencing inhibiting atherosclerotic lesion development and altering plaque composition (Fig. 7d). K_Ca_3.1 silencing leads to alterations in key nuclear factors including downregulation of cFos and NR4A1 and upregulation of PPARα, PPARγ and PGC1α (1.8-fold, p<0.05) subsequently predicted to lead to inhibition of key atherogenic processes including leukocyte infiltration, macrophage activation, SMC proliferation and necrosis, resulting in inhibition of atherosclerosis observed in the current experiment. The expression of PPAR genes identified through RNA-seq DEG analysis, along with *in silico* GO and IPA analysis, for their potential involvement in the mechanisms underlying K_Ca_3.1 knockout-mediated disease protection, were further validated by qRT-PCR (Fig. 7 e-h). Since total BCA samples were used to perform RNA-seq, aorta samples from the corresponding animals were used for validation analysis.

**Figure 7.**
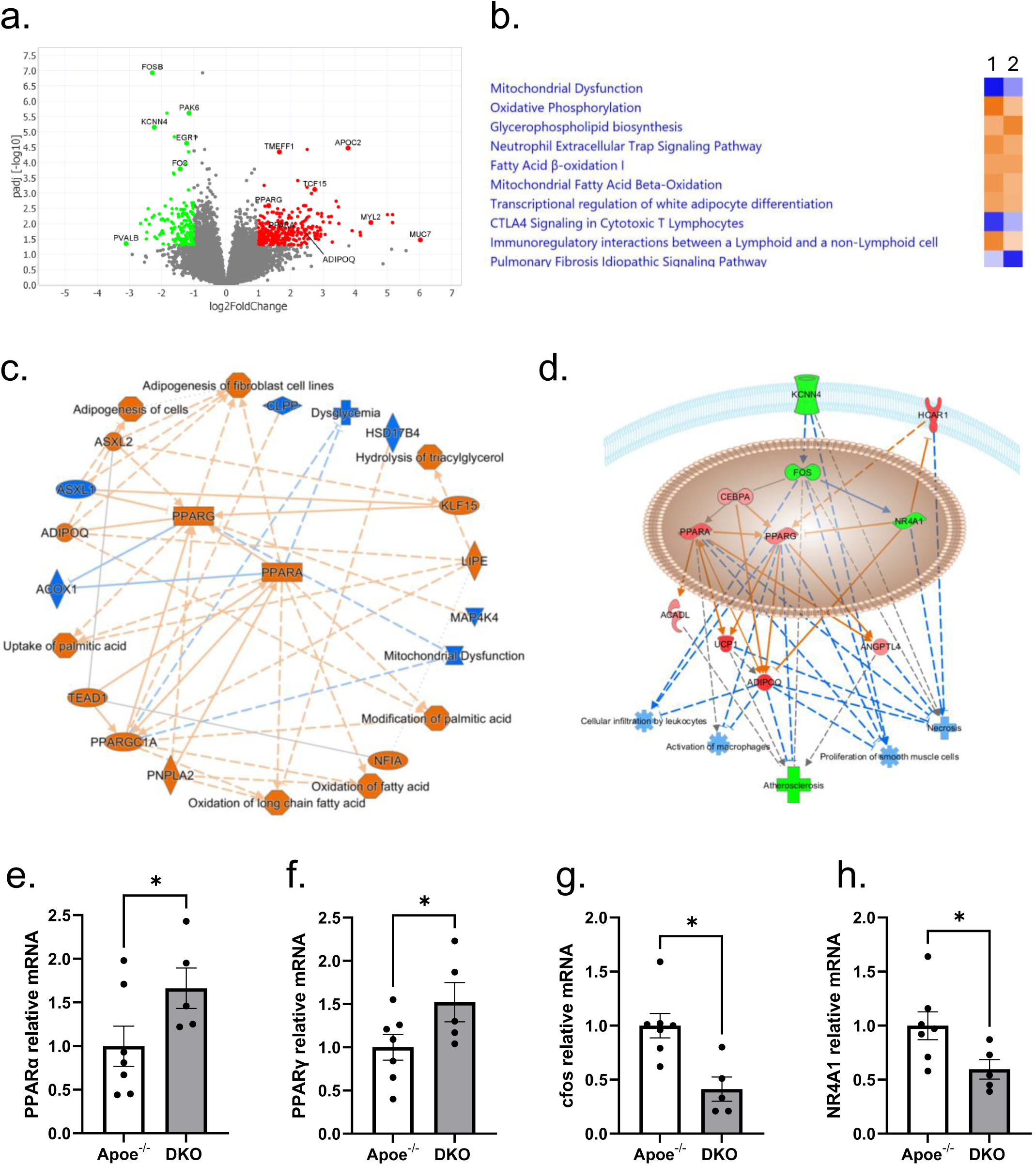
Transcriptomic responses in BCA affected by silencing K_Ca_3.1. (a) Volcano plot of differentially expressed genes in BCA from DKO vs Apoe^-/-^ mice. Upregulated genes coded red and downregulated genes coded green (log^2^fold change <-1 and >1, adjusted p value < 0.05). (b) Comparison of top 10 pathways affected in DKO vs Apoe^-/-^ (1) vs. rosiglitazone treatment of treatment effect on bone marrow mesenchymal stem cells (GSE10192.GPL1261.test6; 2). (c) Graphical summary of effects of K_Ca_3.1 silencing in atherosclerosis as generated by IPA showing the central role of PPAR activation. (d) Downstream regulated biological outcomes based on major DEG genes with of K_Ca_3.1 silencing leading to putative beneficial outcomes related to reduced atherosclerosis, including inhibition of cellular infiltration of leukocytes, reduced activation of macrophages, reduced proliferation of SMCs, and reduced necrosis. Green indicated downregulated genes, red indicates upregulated genes, blue indicates predicted inhibition. Key model genes in d were validated with qRT-PCR in aorta showing significant increases in PPAR (e), PPAR (f), and decreased cFos (g) and NR4A1(h).

## Discussion

The current study is the first to use *in vivo* gene silencing to examine the role of K_Ca_3.1 on atherosclerosis progression. Using a Apoe^-/-^Kcnn4^-/-^ mouse, we were able to demonstrate a reduction in atherosclerotic plaque size, SMC and macrophage content and decreased necrotic core due to silencing K_Ca_3.1. While these outcomes are consistent with previous studies using pharmacological inhibition of K_Ca_3.1 *in vivo*^5,7^, this study provides the first genetic validation of the role of K_Ca_3.1 in regulating atherosclerosis. In addition, subsequent transcriptomic analysis revealed novel putative mechanisms of K_Ca_3.1 silencing of an improved mitochondrial function and a transcriptomic profile similar to that induced by PPAR activation, both of which have been implicated in reducing atherosclerosis.

Our finding of a reduction in SMC content in lesions is consistent with previous studies showing that TRAM-34, an inhibitor of K_Ca_3.1, inhibits smooth muscle cell proliferation and migration^7,8,22^. Accordingly, one would expect a reduction in medial-to-intimal smooth muscle migration and expansion and secondary reduced intimal fibrosis during lesion development with K_Ca_3.1 inhibition. On the contrary, we did not see a reduction in BCA lesion collagen content as previously seen with TRAM-34^12^ and other studies which demonstrated K_Ca_3.1 inhibition reduces fibrosis^10,38–40^. Although the relative collagen content remained unchanged with K_Ca_3.1 silencing, there was a significant reduction in relative necrotic core size in lesions of both male and female DKO mice. Atherosclerotic plaques prone to rupture and subsequent luminal thrombosis are characterized by large necrotic areas, fibrous cap thinning, apoptosis surrounding the necrotic core, and high levels of inflammatory cytokines and matrix proteases^20,41,42^. These thin cap fibroatheromas (TCFA) contribute to major adverse cardiovascular events (MACE), including sudden death. Necrotic cores contain free and esterified cholesterol and dead or senescent macrophages and form due to insufficient efferocytosis of foam cells^43,44^. The significant reduction in necrotic core content of the lesions with K_Ca_3.1 silencing results in lesions with characteristics of a more stable plaque and, consequently, be expected to reduce plaque rupture risk^45,46^. Thus, our study demonstrates that K_Ca_3.1 inhibition not only decreases lesion size but alters plaque composition toward a more stable phenotype.

Moreover, silencing K_Ca_3.1 led to a reduced macrophage content in both male and female lesions as assessed by the macrophage marker, CD68 (Figure 3d-f). This is consistent with previous findings that K_Ca_3.1 inhibitors reduce lesion macrophage content in Apoe^-/-^ mice^7,11^. In macrophages, K_Ca_3.1 has been shown to regulate migration, M1/M2 polarization, respiratory burst and pathogen killing^16^. Pharmacological blockade of K_Ca_3.1 inhibits differentiation toward M1 phenotype and increases M2/M1 ratio in advanced plaques^11^. Inflammation is a potent regulator of collagen deposition and fibrosis, both of which directly influence plaque stability and the severity of atherosclerosis^47^. Inflammatory mediators, such as angiotensin II (Ang II), promote macrophage polarization toward the pro-inflammatory M1 phenotype, which destabilizes plaques by producing enzymes like matrix metalloproteinases (MMPs) that degrade the fibrous cap, weakening plaque integrity^48–50^. In contrast, M2 macrophages, which are anti-inflammatory, help maintain plaque stability by promoting collagen deposition and limiting excessive inflammation, thereby stabilizing the plaque^51,52^. Furthermore, studies have demonstrated that K_Ca_3.1 inhibition reduces vascular inflammation^53^ and may shift macrophage polarization toward the M2 phenotype, thereby contributing to the formation of smaller, more stable plaques^45,46,54,55^. Thus, loss of K_Ca_3.1 in macrophages due to global silencing may contribute to the observed reduction in macrophage infiltration into atherosclerotic lesions, thereby further promoting plaque stabilization.

Alternatively, the findings that both K_Ca_3.1 inhibitors^11^ and gene silencing show reductions in lesion macrophage content could also indicate a role of SMC K_Ca_3.1 through MCP1/Ccl2 expression and/or SMC cell transdifferentiation to foam cells. The cytokine MCP1/Ccl2 is one of several chemokines that play a key role in monocyte recruitment to atherosclerotic plaques^56^. Polymorphisms in the MCP1/Ccl2 promoter thought to increase MCP1/Ccl2 expression have been associated with CAD, chronic stable angina, and myocardial infarction^56^. Using MCP1/Ccl2 receptor-deficient mice to examine atherosclerosis, it was demonstrated that, in the absence of the receptor for MCP1/Ccl2, CCR2, there was a substantial reduction in arterial lipid deposition^57^ and diminished numbers of macrophages in the arterial wall^58^. We provide evidence that K_Ca_3.1 positively regulates MCP1/Ccl2 expression in SMC, but not endothelial cells (Figure 7 d-f). In isolated SMC, inhibition of K_Ca_3.1 by siRNA or TRAM-34 inhibits MCP1/Ccl2 expression, while activation of K_Ca_3.1 with SKA-31 greatly enhances MCP1/Ccl2 expression. Thus, silencing K_Ca_3.1 in SMC *in vivo* may contribute to reduced macrophage recruitment into the intima.

In addition, *in vitro* cholesterol loading of SMCs reduces smooth muscle differentiation markers (e.g. SMαA, SMMHC) and increases macrophage marker expression (CD68, Mac-2)^59,60^. Allahverdian et al. ^61^ concluded that approximately 50% of foam cells in human coronary lesions are of SMC, not monocyte, origin. Similarly, data using SMC lineage tracking in mice have concluded the majority of intimal foam cells (up to 80%) are of SM cell origin^62^. This SMC to foam cell trans differentiation is thought to be associated with a down regulation of myocardin^63^, which we have shown is mediated, in part, by K_Ca_3.1^5,8,22^. Given these observations, the reduced macrophage/foam cell content in lesions following K_Ca_3.1 inhibition may be attributed to effects on SMCs, rather than macrophages infiltration. Thus, K_Ca_3.1 may regulate SMC plasticity and their potential to transition into foam cells, contributing to plaque composition and stability. However, to more precisely define the contributions of SMCs versus macrophages to foam cell formation *in vivo*, future studies utilizing cell-specific knockout and/or lineage tracing models will be essential. These approaches will help delineate the distinct roles of each cell type in lesion formation and foam cell accumulation, ultimately advancing our understanding of the molecular mechanisms driving atherosclerosis.

In addition to the potential mechanisms underlying the beneficial effects of K_Ca_3.1 inhibition on atherosclerosis, transcriptomic analysis revealed novel effector signaling pathways associated with K_Ca_3.1. Specifically, this analysis highlighted a significant enhancement of mitochondrial function and the activation of PPAR signaling upon K_Ca_3.1 silencing. Pathway analysis using Ingenuity Pathway Analysis (IPA) showed a striking similarity between the top canonical pathways affected in BCAs following K_Ca_3.1 silencing and those influenced by the PPAR agonist, rosiglitazone (Figure 6b, c). These pathways included improved mitochondrial function, enhanced fatty acid oxidation, and the inhibition of fibrosis, all of which are known to contribute to atherosclerosis regression and plaque stabilization. Previous studies have shown that PPAR agonists reduce vascular inflammation, plaque size, and atherosclerosis progression^64–67^, as well as lower in-stent NI volume in non-diabetic patients ^68^. For example, rosiglitazone has been shown to decrease atherosclerotic plaque size by reducing lipid deposition and macrophage infiltration, while also decreasing CCL2/Mcp1 levels and increasing adiponectin in the aorta ^64^ all similar to effects observed in the current study. However, given that endothelial cells, vascular smooth muscle cells (SMCs), monocytes/macrophages, and T cells all express PPARα and PPARγ ^69^, it is difficult to pinpoint which specific cell types are primarily responsible for the beneficial effects observed. To address this, further cell-specific studies, including lineage tracing and knockout models, will be essential to delineate the exact cellular mechanisms underlying the improvements in plaque stability and regression associated with K_Ca_3.1 inhibition.

Increased mitochondrial respiration and reduced mitochondrial dysfunction have been consistently associated with decreased atherosclerotic progression^70–72^. In the aortas of Ob/Ob, LDLR^-/-^ mice, low levels of cytochrome oxidase were linked to increased plaque formation and elevated MCP1/Ccl2 levels^70^. Similarly, in pig coronary plaques, cytochrome oxidase I and 4I1 were found to be reduced in macrophages from complex Stary III plaques compared to less advanced Stary I plaques^70^. Hypercholesterolemia results in a coordinated down-regulation of mitochondrial genes. Moreover, these genes are tightly connected in co-expression network modules related to mitochondrial biogenesis and antioxidant responses, possibly regulated by ERR-α/PGC1-α^73^. These findings suggesting that improved mitochondrial activity could contribute to the stabilization and regression of atherosclerotic plaques. The effects of K_Ca_3.1 inhibition on mitochondrial function observed in our study align with these previous findings. Our transcriptomic analysis reveals that silencing K_Ca_3.1 not only enhances mitochondrial function but also activates signaling pathways similar to those triggered by PPAR agonists, such as rosiglitazone. Notably, rosiglitazone has been shown to increase mitochondrial biogenesis and function in tissues like the brain^74^ and adipose tissue^75^, and these processes are known to be protective against atherosclerosis. The parallels between the effects of rosiglitazone and the transcriptomic changes induced by K_Ca_3.1 silencing suggest that activation of the PPAR pathway may play a central role in the beneficial effects of K_Ca_3.1 inhibition. Notably, K_Ca_3.1 silencing increased UCP1 in BCAs in the current study. UCP1 is activated by PPARα and PPARγ^76^ has been shown to inhibit atherosclerosis by reducing vascular inflammation^77^. This connection further underscores the potential for targeting K_Ca_3.1 as a therapeutic strategy for atherosclerosis, as improving mitochondrial function and activating PPAR pathways could collectively reduce vascular inflammation, improve plaque stability, and limit disease progression. Additionally, the observed effects on mitochondrial function and PPAR signaling provide critical mechanistic insights into how K_Ca_3.1 silencing may promote atherosclerosis regression and plaque stabilization.

In conclusion, our study provides further support for the important contribution of K_Ca_3.1 activation in the progression of atherosclerotic lesion development and composition and provides novel insights into mechanisms of the beneficial effect of K_Ca_3.1 inhibition on atherosclerosis. Together, our *in vivo*, *in vitro* and transcriptomic findings provide a novel model of the mechanisms underlying K_Ca_3.1 silencing inhibiting atherosclerotic lesion development and altering plaque composition (Fig. 7d) involving key nuclear factors cFos, NR4A1, PPARα, PPARγ and PGC1α leading to reduction in plaque size and altered composition. Together with previous studies, these finding support the potential therapeutic application of pharmacological inhibition of K_Ca_3.1 in limiting the progression of atherosclerosis. However, there are limitations to the current study. As noted, increased expression of the K_Ca_3.1 has been observed in multiple cell types involved in atherosclerotic lesion progression including smooth muscle, macrophages and T cells. Studies to date, including the current study, have relied on global gene silencing or systemic pharmacological interventions *in vivo*. Similarly, the relative contribution of K_Ca_3.1 among cell types has not been determined and will require cell-specific, inducible silencing/overexpression and, potentially, fate tracking to elucidate responsible cell type(s).

## Conflict of interest

None.

---

***Supplementary Figure 1:*** Effects of K_Ca_3.1 inhibition and activation on Col1a1 expression. (a,b) Neither inhibition of KCa3.1 with siRNA nor treatment with TRAM-34 affected Col1a1 expression. (c) However, activation of KCa3.1 increased Col1a1 expression, independent of PDGF-BB. Data represented as mean ± SE. the *p*-value ≤0.05 was considered statistically significant. * *p*-value ≤0.05. # *p*-value >0.05.

***Supplementary Figure 2:*** *The metascape analysis:* (a) Bar graph showing enriched terms across input gene lists and (b) output of regulatory factor analysis, both colored by *P*-values.

**Supplementary Table 1:**
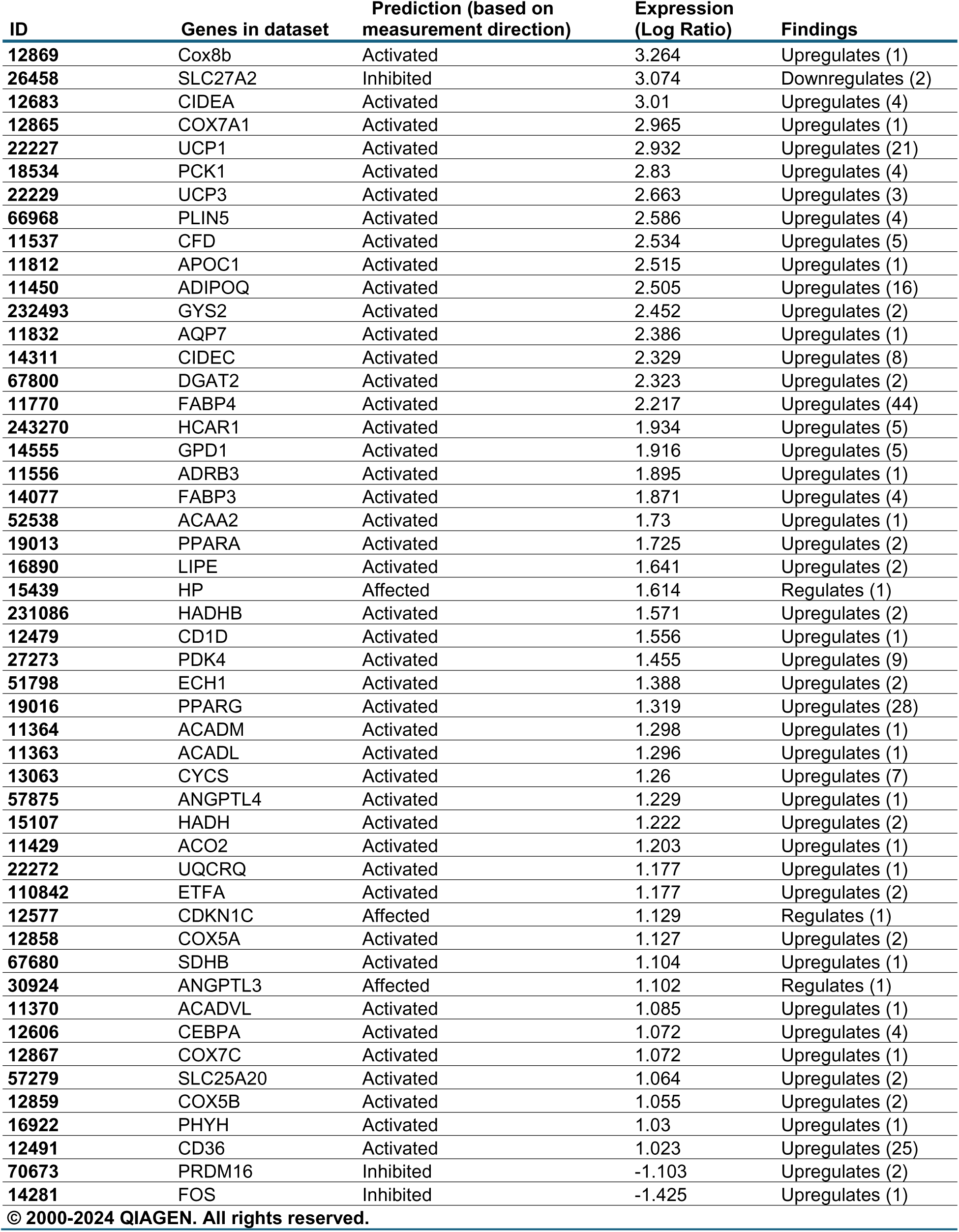
Up and down regulated genes in DKO associated with the IPA-identified top upstream regulator, i.e. the PPAR agonist, rosiglitazone. DKO resulting in differential expression in 45 of 52 predicted DEGs associated with rosiglitazone.

## Notes

### Competing Interest Statement

The authors have declared no competing interest.

