## Supplementary figures and images for "Genetic silencing of K_Ca_3.1 inhibits atherosclerosis in ApoE null mice"

### Supplementary Fig. 1

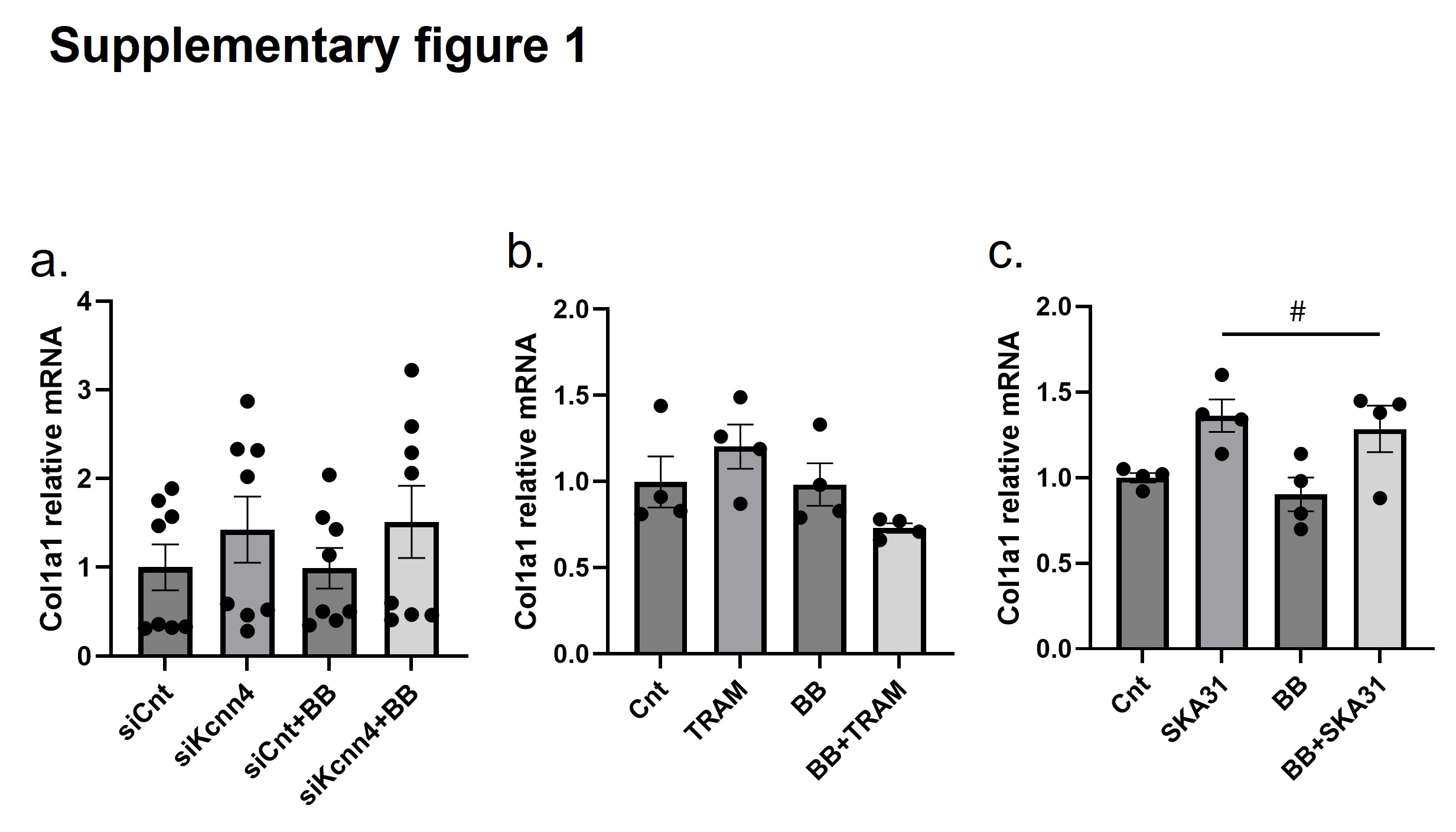

### Supplementary Fig. 2

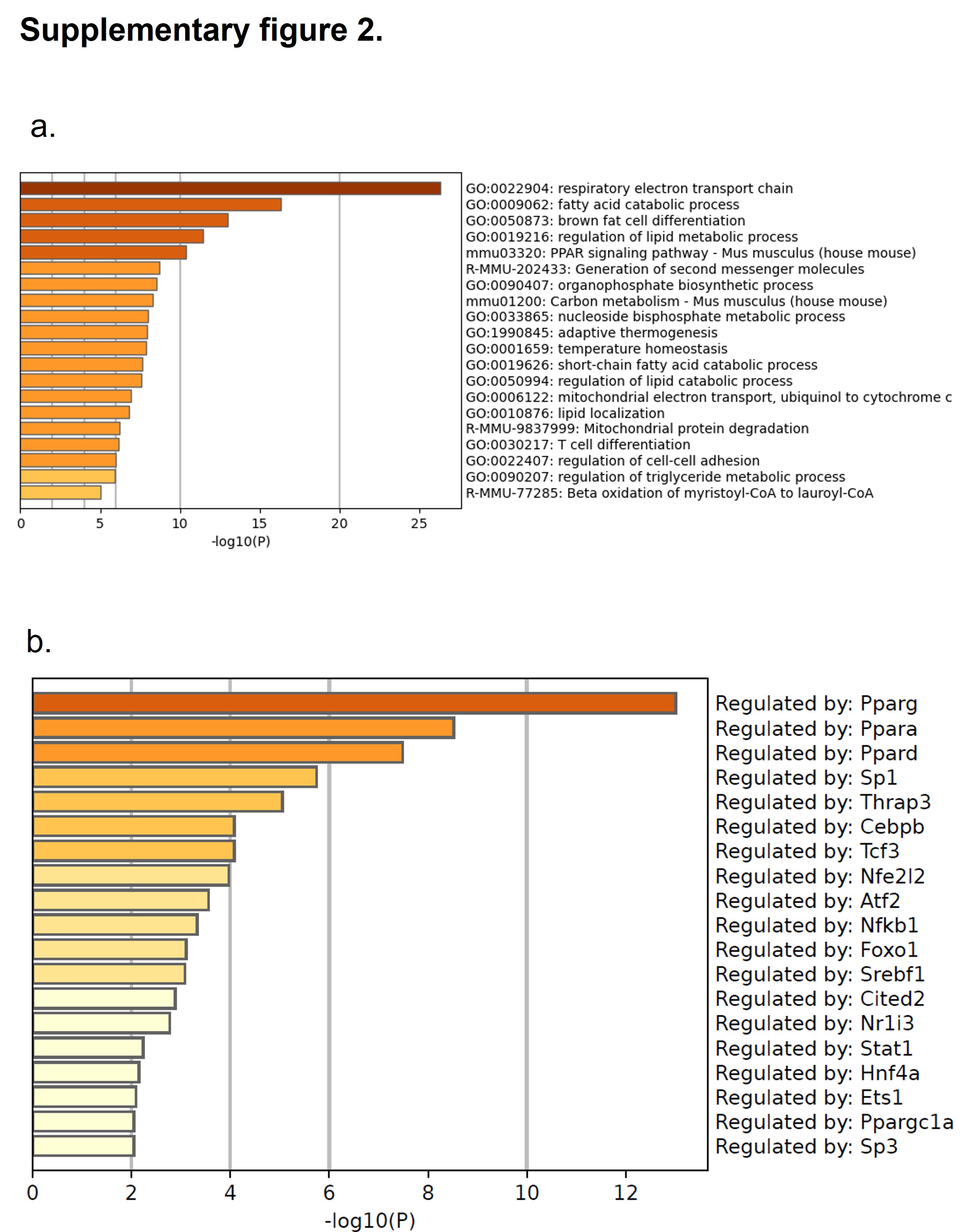
